# The piRNA pathway sustains adult neurogenesis by reducing protein synthesis and cellular senescence

**DOI:** 10.1101/2020.09.15.297739

**Authors:** C. Gasperini, K. Tuntevski, R. Pelizzoli, A. Lo Van, D. Mangoni, R.M. Cossu, G. Pascarella, P. Bianchini, P. Bielefeld, M. Scarpato, M. Pons-Espinal, R. Sanges, A. Diaspro, C.P. Fitzsimons, P. Carninci, S. Gustincich, D De Pietri Tonelli

## Abstract

Adult Neural progenitor cells (aNPCs) ensure lifelong neurogenesis in the mammalian hippocampus. Proper regulation of aNPC fate entails important implications for brain plasticity and healthy aging. Piwi proteins and the small noncoding RNAs interacting with them (piRNAs) are best known in gonads as repressors of transposons. Here, we show that Piwil2 (Mili) and piRNAs are abundant in aNPCs of the postnatal mouse hippocampus and demonstrate that this pathway is essential for proper neurogenesis. Particularly, depleting the piRNA pathway in aNPCs impaired neurogenesis, increased senescence and accordingly the generation of reactive glia. Moreover, this manipulation primarily elevated 5S ribosomal RNA, SINEB1 and mRNAs encoding ribosomal proteins and regulators of translation, resulting in higher polysome density and protein synthesis upon differentiation. Our results provide evidence of an essential role for the piRNA pathway in maintaining homeostasis to sustain neural stem cell fate, underpinning its possible involvement in brain plasticity and successful aging.

## Introduction

A regulated balance of neural progenitor cells’ (aNPCs) quiescence, proliferation and differentiation guarantees lifelong neurogenesis in the adult hippocampus (Altman, 1962; Doetsch et al., 1999), prevents the generation of reactive glia (Encinas et al., 2011; clarke 2018; Sierra et al., 2015), and neurodegeneration (Toda et al., 2019). Understanding the molecular control of aNPCs fate is pivotal to develop novel therapies to prevent or delay age dependent-loss of neurogenesis and related pathological conditions.

Piwi belong to an evolutionary conserved subfamily of Argonaute proteins that bind to Piwi-interacting RNAs (piRNAs), single-stranded noncoding RNAs of 21-35 nucleotides. These Piwi proteins and piRNAs (henceforth referred to as the piRNA pathway) are highly abundant in gonads, where they mainly target transposable elements (TEs) for degradation to maintain germline stem cell pools and male fertility (Czech et al., 2018; Ozata et al., 2019).

Since its initial discovery, the piRNA pathway has been implicated in regulating gene expression outside of the germline. Moreover, it is preferentially expressed in germ cells, embryonic stem cells and adult stem cells, compared to their differentiated progeny (Rojas-Ríos and Simonelig, 2018). Indeed, expression of Piwi proteins was observed in human hematopoietic stem cells, suggesting a possible role of this pathway in maintaining the stem cell pool (Sharma et al., 2001), whereas its function is dispensable for normal hematopoiesis in mouse (Nolde et al., 2013).

In the adult brain, the piRNA pathway was proposed to control synaptic plasticity and memory (Lee et al., 2011; Leighton et al., 2019; Nandi et al., 2016; Zhao et al., 2015). PiRNA levels in neurons, however, are low compared to germline cells (Lee et al., 2011; Nandi et al., 2016), disputing the proposed piRNA pathway function in the nervous system. Apart from gonads, the highest piRNA expression in adult mice has been found in the hippocampus (Perera et al., 2019), however, the specific cell type in which piRNA expression takes place was not identified. Concomitantly, transposable element (TE) expression is high in differentiating NPCs (Muotri et al., 2005), and Piwil1 (Hiwi) was recently shown to regulate human glioma stem cell maintenance (Huang et al., 2021). Thereby, it is reasonable to hypothesize that the piRNA pathway may play a role in aNPCs maintenance and/or differentiation as well.

Here, we demonstrate that the piRNA pathway is enriched in aNPCs, and that its function is essential for proper neurogenesis. Moreover, we show that inhibiting this pathway in aNPCs leads to senescence-associated inflammation in the postnatal mouse hippocampus, a condition which has been linked to age-related impairment of neurogenesis and increase in astrogliogenesis (Clarke et al., 2018; Encinas et al., 2011). Our results implicate the piRNA pathway in the aging of the hippocampal stem cell niche.

## Results

### Mili expression is abundant in aNPCs and depleted in neurogenesis

As an entry point to investigate Piwi pathway in aNPCs we quantified expression of *Piwil1* (Miwi) *Piwil2* (Mili) and *Piwil4* (Miwi2) mRNAs, which encode the essential proteins for piRNA biogenesis and function in mammals (Czech et al., 2018; Ozata et al., 2019), in cultured aNPCs derived from neural stem cells (NSC) of the adult mouse Dentate Gyrus (DG) (Babu et al., 2011; Pons-Espinal et al., 2017), by RNA sequencing (RNA seq) (Fig.1A). Of the three main Piwi genes in mouse, Mili was the most abundant in aNPCs, and its expression increased transiently upon induction of their spontaneous differentiation. The Mili transcript was particularly more abundant at day 4 (here referred to as days of differentiation DIF 4) upon onset of spontaneous differentiation compared to undifferentiated (i.e., proliferating) aNPCs (DIF 0), whereas it was depleted in differentiated cells (DIF 7-14) (Fig 1A). Similarly, the Mili protein abundance (Fig. 1B) was higher in undifferentiated aNPCs (DIF 0) or early upon onset of vector-induced neurogenesis (DIF 4, neuroblasts) than in differentiated neurons (DIF 7-14). Next, we quantified the abundance of Miwi and Mili proteins in the mouse testis, whole hippocampus, and compared it to the one in undifferentiated aNPCs (Fig. 1C, D). As expected, the Miwi protein was very abundant in testis, but was not detectable in the whole hippocampus or aNPCs (Fig.1C), whereas the Mili protein abundance in aNPCs was about 40% of the one in the testes (Fig. 1D), and about four-fold higher than in primary hippocampal neurons (Fig. 1E). To validate this finding *in vivo*, we used a previously published split-Cre viral approach to selectively label NSCs and their progeny in the hippocampus of postnatal Td-Tomato Cre-reporter mice (Beckervordersandforth et al., 2014; Pons-Espinal et al., 2017). Five days post viral-injection (dpi), we found Mili protein presence in Td-Tomato positive (Td+) NSCs of the subgranular zone (SGZ) of the DG (Fig. 1F). To quantify *Mili* during neurogenesis *in vivo*, we sorted Td+ NSCs and their differentiated progeny at 10 and 30 dpi in the postnatal mouse hippocampus, respectively. The *Mili* transcript was significantly more abundant in Td+ NSCs (10 dpi) than in adult-born Td+ neurons (30dpi) or Td-cells (Fig. 1G). These results demonstrate that Mili expression is enriched in neural stem/progenitor cells and depleted in their progeny.

**Fig. 1.**
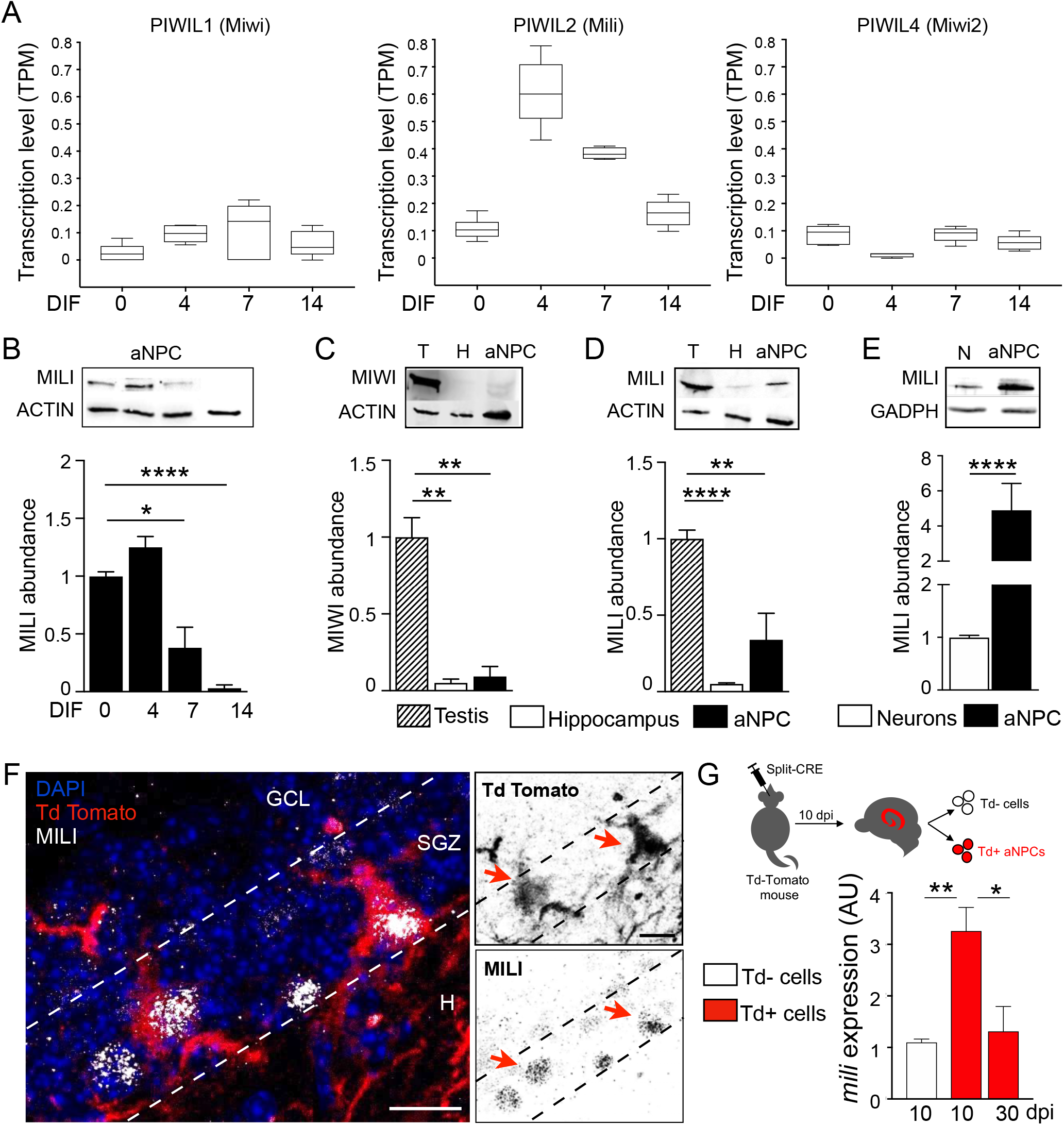
Hippocampal expression of Mili is enriched in aNPCs and decreases during neurogenesis. (A) Levels of PIWIL1 (Miwi), PIWIL2 (Mili) and PIWIL3 (Miwi2) transcripts in RNA seq. data from aNPCs cultures in proliferative (Days of differentiation –DIF– 0) and differentiation media (DIF 4 to 14). (B) Mili protein abundance in undifferentiated aNPCs (DIF0) and upon viral-induced neurogenesis (DIF4-14). (C) Miwi and (D) Mili protein abundance in extracts from postnatal mouse testis, hippocampus and from undifferentiated aNPCs cultures. (E) Mili protein abundance in in extracts from cultured mouse hippocampal neurons and undifferentiated aNPCs. (F) Mili (white) expression in Td+ NSC (red) in hippocampal subgranular zone (SGZ); arrows indicate Td+ Mili + double-positive cells. (G) Scheme of the experiment (top), *Mili* mRNA expression in sorted Td+ and Td– cells after *in vivo* transduction with split-Cre viruses in the hippocampus (bottom). Data are expressed as mean ± SEM, n = 3 independent experiments. t-Student test or one-way ANOVA Bonferroni as post hoc: *p < 0.05, **p < 0.01, **** p < 0.0001. GCL, granular cell layer; H, Hilus. Scale bars: 10 µm.

### Level of piRNAs parallels Mili abundance in neurogenesis

We next wished to determine whether piRNAs are found in aNPCs and quantify their abundance in neurogenesis by small RNA seq (Fig. 2). Following a previously published analysis pipeline (Ghosheh et al., 2016), we identified a total of 725,472 *bona fide* piRNAs, of which 33,396 perfectly aligned (*i*.*e*, no mismatch) with those previously annotated in the piRNA database (piRBase (Zhang et al., 2014)), had an average length of 30 nt (Fig. 2A) and bore a 5’ uridine (U) bias (Fig. 2B). Moreover, the distance probability between the 5’ termini of putative primary and secondary piRNAs was distributed similarly to that of other animals (Gainetdinov et al., 2018), with asymptotic convergence around the ‘0’ mark on the abscissa (Fig. 2C). Mature piRNAs typically bear 2’-*O*-methylation at their 3’ termini, which confers them stability and enables stronger binding to Piwi proteins (Czech et al., 2018; Ozata et al., 2019). We tested whether the putative piRNAs isolated from aNPCs were also methylated by evaluating their resistance to periodate oxidation and alkaline ß-elimination, as previously reported (Kirino and Mourelatos, 2007). Indeed, a quantitative real time PCR (qPCR)-based small RNA assay (Taq-Man) indicated that some of the most abundant piRNAs expressed in aNPCs exhibited resistance to periodate treatment, thus indicating their methylation (Fig 2D). In contrast, control synthetic RNAs or endogenous small noncoding RNAs, such as snoRNA bearing a terminal 2’,3’-hydroxyl group, were oxidized after the treatment and therefore degraded (Fig 2D). Furthermore, to assess whether putative piRNAs were depending on Mili, we quantified their expression in aNPC cultures upon *Mili* knockdown (Mili-KD) achieved by transducing short-hairpin RNAs targeting *Mili* transcripts, or a scramble short-hairpin RNA as control (or two different synthetic GapmeRs targeting *Mili*, Fig. S1), through a viral vector-mediated protocol *in vitro*. Indeed, reduction of Mili protein abundance in aNPCs (Fig.2E) was sufficient to deplete four of the most highly expressed piRNAs in aNPCs (Fig. 2F and Fig. S1), in agreement with the conclusion that Mili is the most abundant Piwi protein in these cells (Fig.1). Of note, this manipulation did not affect Miwi expression, excluding possible compensatory effects (Fig. S1). Together, this evidence indicates that the most abundant sequences homologous to known piRNAs found in aNPCs fulfill at least five of the criteria that characterize *bona fide* piRNAs (*i*.*e*., length, U-Bias, 2’-O-Methylation at the 3’ end, inter-distance and MILI dependence).

**Fig. 2.**
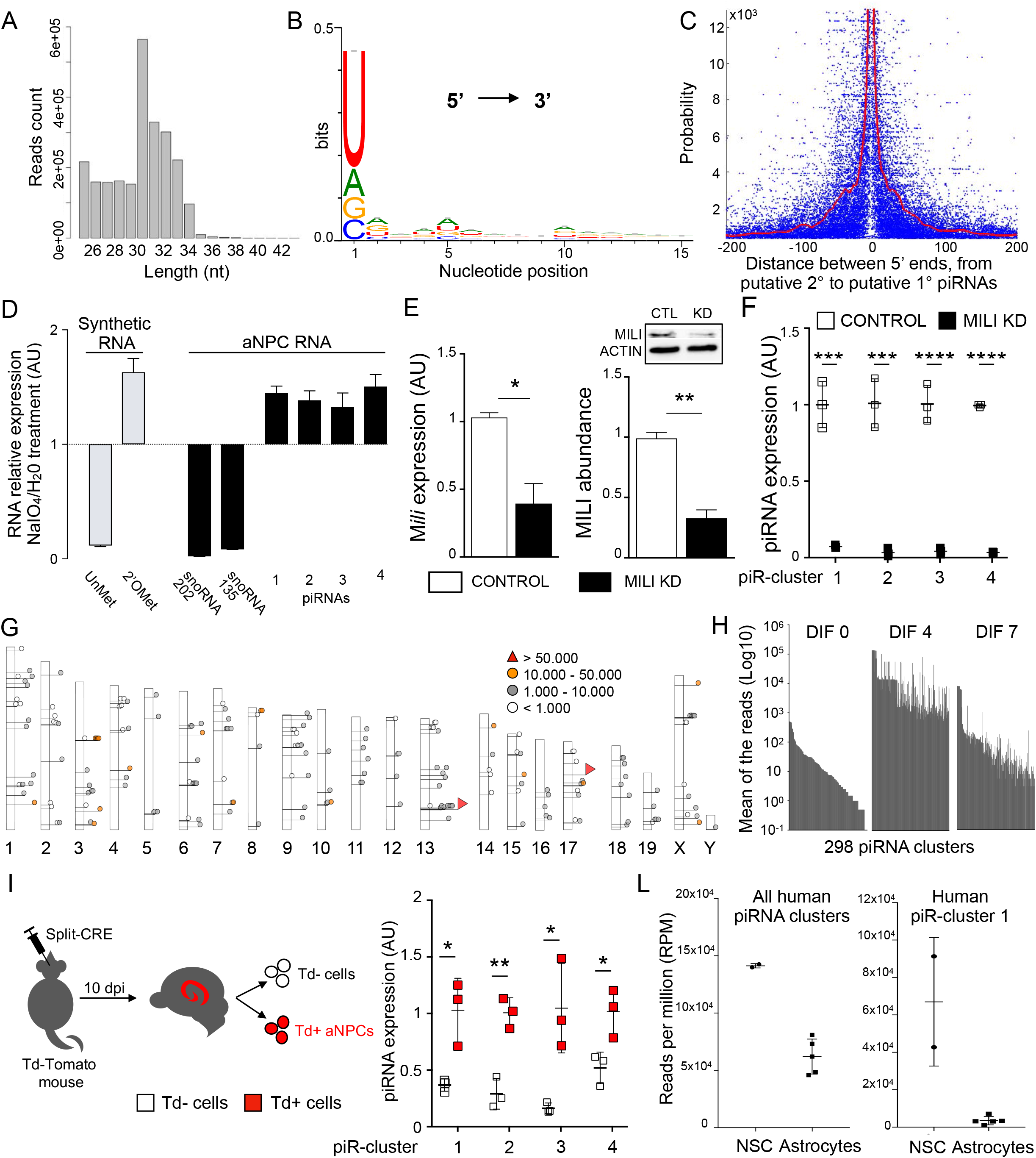
Level of piRNAs parallels Mili abundance in neurogenesis. (A) Size distribution of piRNA reads. (B) Uridine bias at piRNA 5’ ends. (C) Probability of distances from the 5’ends of putative secondary piRNAs to the 5’ends of putative primary piRNAs. Note that 5’ termini of putative primary and secondary piRNAs derived from the same cluster tend to concatenate around the ‘0’ mark, as reported in other animals. Distance probability was assayed for unique piRNAs (length between 15-35 nucleotides), without taking into account abundance, by locally weighted smoothing linear regression (LOWESS). (D) RNA relative expression of four of the top abundant piRNAs (in H) and two control snoRNAs (202 and 135) in aNPC, upon treatment with sodium periodate (NaIO4) or water and alkaline ß-elimination. Synthetic RNA oligos were used as negative (Unmethylated, UnMet) and positive (2’-O-methylated, 2’OMet) controls, respectively. Note that presence of 3’-end 2’-O-methylation in positive control and piRNAs confers them resistance to periodate oxidation and alkaline ß-elimination, in contrast to their depletion in UnMet negative control and snoRNAs. (E) *Mili* mRNA (left) and protein (right) levels in aNPCs upon viral transduction of scrambled shRNA (Control) or shRNA targeting *Mili* (Mili KD). (F) Expression of four of the most abundant piRNA clusters in control and Mili KD aNPCs. (G) Genomic location of 298 piRNA clusters on mm9. (H) Mean reads of piRNA clusters in aNPCs (DIF0) and upon viral-induced neurogenesis (DIF4-7), where the total number of filtered reads in each library ranges between 1,4 -4,4 × 10^6^. (I) (left) Schematic representation of the experiment; (right) expression of four of the most abundant piRNA clusters in sorted Td+ and Td-cells 10 dpi of split-Cre viruses in hippocampus. (L) Expression of piRNA clusters (left) and piR-cluster 1 (right) in human NSC and astrocytes. Data are expressed as mean ± SEM, n = 2 (A-C, G-H) and n = 3 (D-F, I) independent experiments. t-Student test as post hoc: *p < 0.05, **p < 0.01, *** p < 0.001, **** p < 0.0001.

Genomic mapping of the piRNA reads from aNPCs and their progeny identified 298 clusters perfectly aligning to the mouse genome (Fig 2G and Table S1), with an average size of 168 bases and some of them exceeding 2000 bases, as seen in mouse testes (Aravin et al., 2006; Girard et al., 2006). The piRNA raw reads/cluster averaged around 4700 reads, with two clusters, one located in the chromosome 13 and one in the 17, giving rise to more than 80,000 piRNA reads (Fig. 2G). Analysis of small RNA seq data for directionality suggested a strand bias where the majority of the piRNAs arise unidirectionally, although bidirectional piRNAs were also identified. Analysis of piRNA transcript levels in neurogenesis indicated a transient peak at the onset of differentiation (Fig. 2H DIF 4 and Fig. S1), in agreement with the Mili expression pattern. Next, we validated four of the most abundant piRNA clusters in Td+ NSCs sorted from the adult hippocampus confirming their expression *in vivo* (Fig. 2I). Interestingly, one of the piRNA-clusters in our dataset (i.e., piR-cluster 1) is homologous to the human piR-61648 that was recently shown to be selectively expressed in somatic tissues, but is depleted in gonads (Torres et al., 2019) Fig. S1 and Table S1). This observation prompted us to extend our analysis to human NSCs. Analysis of small RNA datasets from the RIKEN FANTOM5 project (De Rie et al., 2017) confirmed the enriched expression of piR-cluster 1, as well as of many additional piRNA clusters in human NSCs compared to differentiated brain cells (Fig. 2L and Table S2). Together, these results indicate that piRNAs are selectively enriched in both mouse and human neural stem/progenitor cells, thus matching the expression of Mili (Fig.1).

### The piRNA pathway sustains proper neurogenesis

To infer functions of the piRNA pathway in aNPCs differentiation, we knocked down (KD) *Mili* by injecting a synthetic antisense oligonucleotide (GapmeR, MILI KD), or a scrambled GapmeR (Control), in the postnatal mouse hippocampus (Fig. 3A). Mili-KD (Fig. 3B) did not alter stemness or growth of aNPCs *in vitro* (Fig. S2) but led to a dramatic increase in the expression of the glial cell marker glial fibrillary acidic protein (GFAP) as early as 48 hours after GapmeR injection *in vivo* (Fig.3B) and *in vitro* (Fig. S2). Inspection of brain sections 30 days after bilateral injections indicated a marked increase in GFAP+ cells with enlarged somas in the ipsilateral hippocampus injected with GapmeR antisense to *Mili*, compared to cells in the contralateral side injected with control GapmeR (Fig. 3C). To ascertain whether GFAP+ cells were actively generated upon Mili-KD, we administered bromodeoxyuridine (BrdU) immediately after GapmeRs injection to label dividing cells in a third cohort of mice (Fig. 3A). 30 days after GapmeR injection, we found that Mili-KD led to a significant increase in adult born GFAP+BrdU+ glial cells at the expense of NeuN+BrdU+ neurons (Fig. 3D). This result was corroborated by gene set analysis in RNA seq data from differentiating aNPCs *in vitro*, showing an enrichment in the expression of astrocyte-related genes and a concomitant deregulation of neuronal fate genes upon piRNA pathway depletion (Fig S2). These results indicated that the piRNA pathway is required to sustain neurogenesis in the postnatal hippocampus at the expense of gliogenesis.

**Fig. 3.**
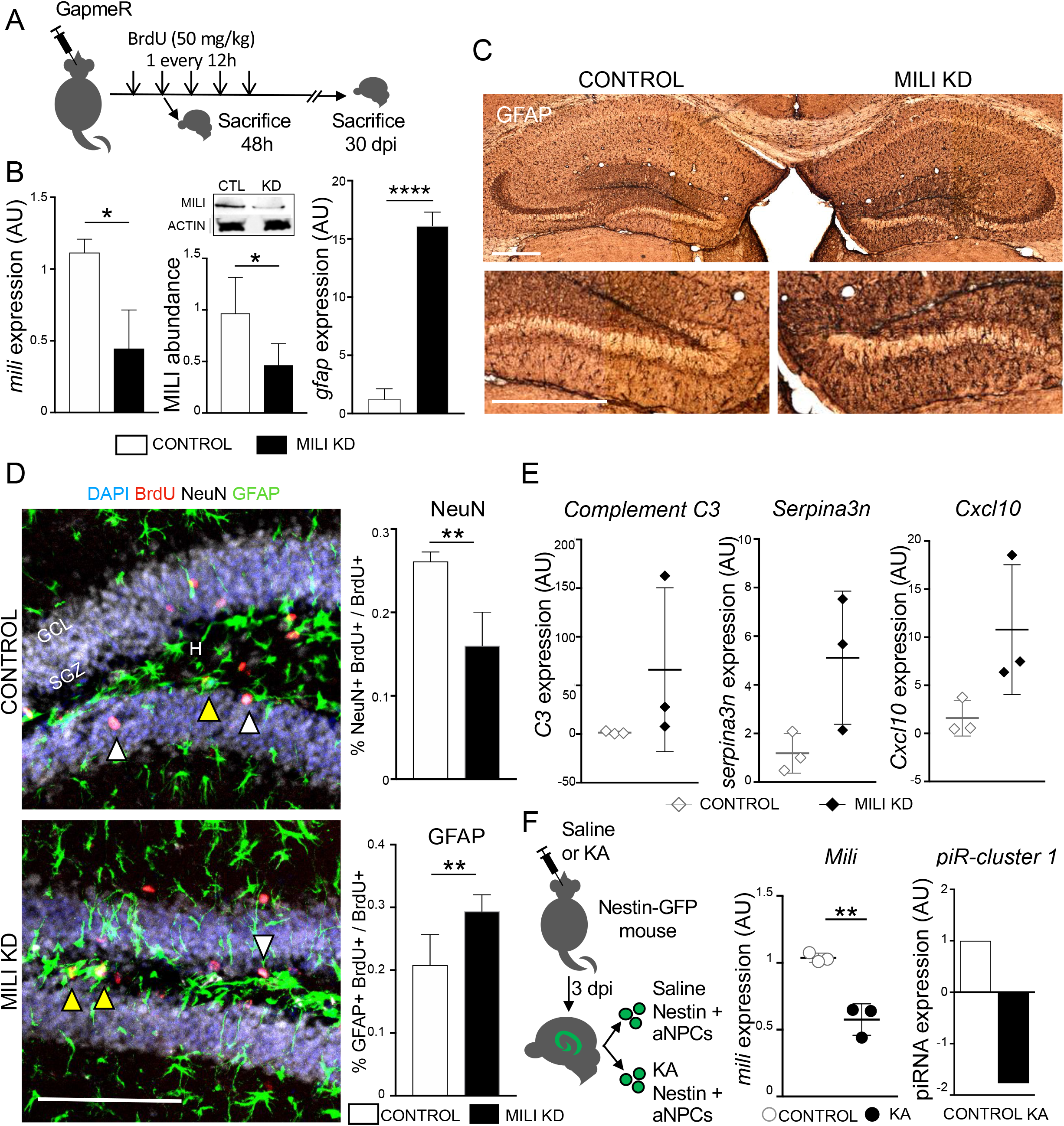
The piRNA pathway sustains proper neurogenesis. (A) Scheme of the *in vivo* experiment. (B) *Mili* mRNA (left), protein (middle) and *Gfap* mRNA (right) levels in the hippocampus 48 hours after the injection of scrambled (Control) or GapmeR against Mili (Mili KD). (C) Light-microscopy images of postnatal hippocampal sections, immunostained for GFAP, 30 dpi of scrambled (Control, left hemisphere) and GapmeR against Mili (Mili KD, right hemisphere). (D) (Left) Immunostaining for GFAP (green), BrdU (red), NeuN (white) and nuclear DNA (blue) in the hippocampus 30 dpi of scrambled (Control) or GapmeR against Mili (Mili KD); (right) percentages of NeuN+BrdU+ (white arrowheads), or GFAP+BrdU+ (yellow arrowheads) over total BrdU+ cells. (E) mRNA expression of reactive astrocyte markers in the hippocampus 48h upon injection of scrambled (control) or GapmeR against Mili (Mili KD). (F) *Mili* mRNA (left) and piR-cluster 1 (right) expression in sorted GFP+ NSCs from Nestin-GFP mice treated with Saline (Control) or Kainic Acid (KA). Data are expressed as mean ± SEM, n = 3 independent experiments (A-C); n=5 (E); n = 7 (F, G); n = 3 (H, I). t-Student test as post hoc: *p < 0.05, **p < 0.01, **** p < 0.0001. Scale bars: 1 mm (C); 100 μm (D).

Increased GFAP expression is generally regarded as a hallmark of astrocyte reactivity (Escartin et al., 2021). In agreement, we observed a significant increase in the levels of known reactive glial markers (Clarke et al., 2018; Liddelow et al., 2017) upon Mili KD in the postnatal hippocampus (Fig. 3E). To confirm this result, we took advantage of Kainic Acid injection in the postnatal hippocampus of mice expressing GFP under the control of the NSCs/NPCs specific promoter *Nestin* (Fig. 3F), a treatment previously shown to induce aNSC conversion into reactive glia (Bielefeld et al., 2017; Sierra et al., 2015). Indeed, this treatment reduced levels of the piRNA pathway in sorted *Nestin-*GFP+ NSCs (Fig. 3F). Altogether, these results demonstrate an essential role of the piRNA pathway for proper neurogenesis and suggest that its downregulation mediates reactive gliogenesis in the postnatal mouse hippocampus.

### Inhibition of the piRNA pathway in aNPCs results in senescence-associated phenotypes

Conversion of hippocampal NSC into reactive glia, at the expense of neurogenesis, has been observed in normal aging (Clarke et al., 2018), and it has been related to increased neuroinflammation and cellular senescence (Babcock et al., 2021; Martín-Suárez et al., 2019). Whether the piRNA pathway is involved in this mechanism in the brain, is unknown. This prompted us to analyze RNA seq data for senescence-associated secretory phenotype (SASP) in Mili-KD in neuroblasts (Fig. 4A). Interestingly, we found that depletion of the piRNA pathway led to a significant increase in the expression of several immune-modulatory and senescence associated genes in neuroblasts compared to control cells (Fig. 4A). Moreover, after piRNA pathway depletion we observed a higher proportion of cells positive for senescence-associated β-galactosidase (β-gal) as early as 48 hours upon induction of their spontaneous differentiation (Fig. 4B). At the same time point, we immuno-stained aNPCs with anti-KI67 (a protein that is expressed in all phases of the cell cycle except G0 and early G1 (Yu, 1992)) and anti-BrdU antibodies. Quantification of the BrdU + and KI67-cells over total BrdU+ indicated a premature cycle exit upon piRNA pathway depletion (Fig. 4C). Similarly, by propidium iodide incorporation and flow cytometry analysis we found an increase in the proportion of cells in G0/G1 phase and a concomitant decrease in S phase cells (Fig. 4D). Accordingly, piRNA pathway depletion did not lead to apoptosis (Fig. S3), whereas we found that it led to altered expression of genes previously associated with senescence-induced cell cycle exit (Fig. 4E, F), circadian mechanism and oxidative stress (Fig. S3), as previously reported (Adusumilli et al., 2021; Babcock et al., 2021; Schouten et al., 2020). To corroborate this evidence *in vivo*, we quantified the expression of *Mili* transcript in sorted *Nestin*-GFP+ NSCs from the DG of young (6 Weeks) and ∼12 Months old mice (i.e., when the majority of hippocampal aNSCs turn into an aged phenotype (Martín-Suárez et al., 2019)) and found that *Mili* transcript was reduced in aged aNSCs (Fig. 4G). These results suggest that the piRNA pathway prevents aNPCs/neuroblasts senescence.

**Fig. 4.**
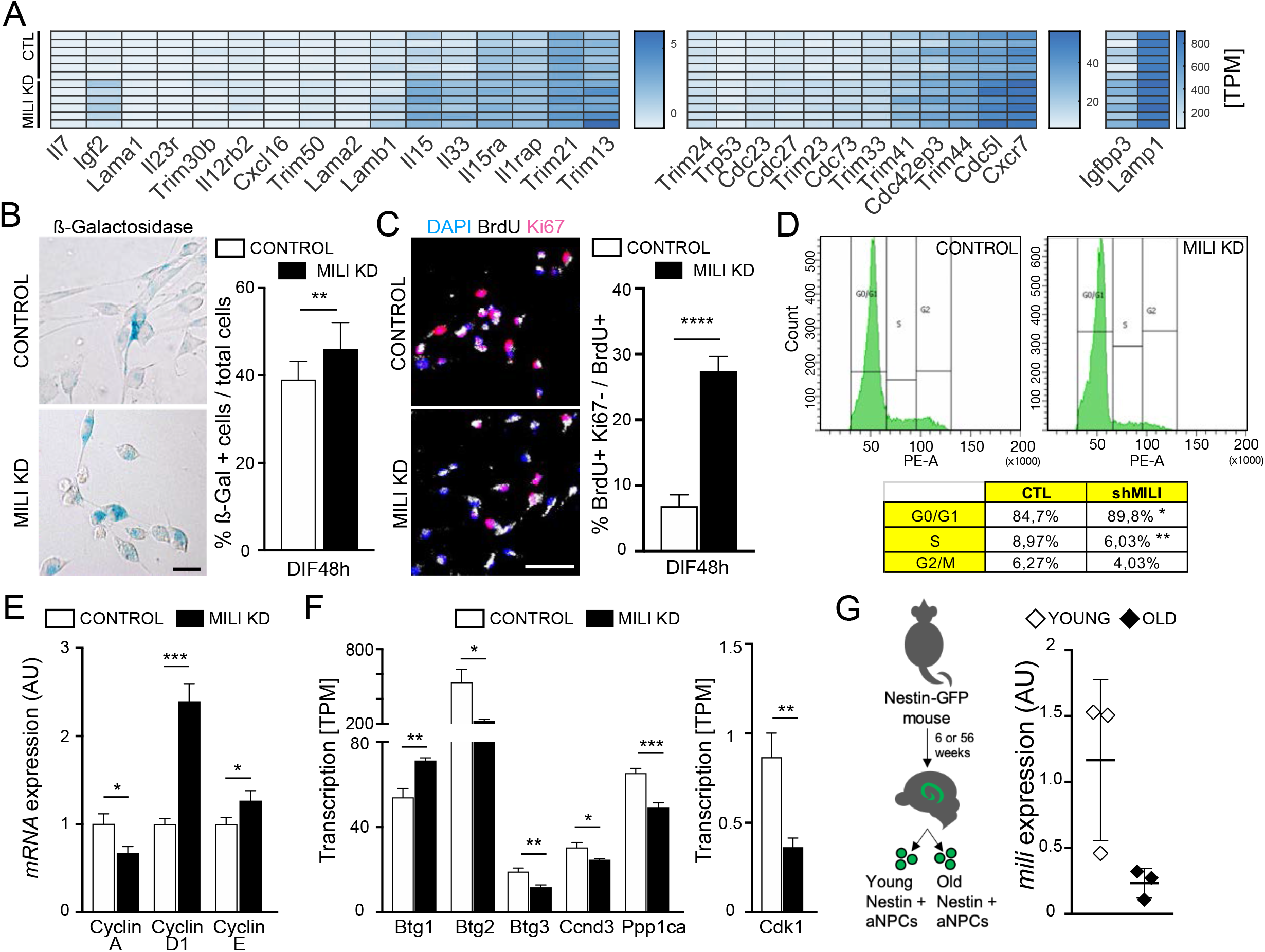
Inhibition of the piRNA pathway in aNPCs induces senescence-associated phenotypes. (A) Heatmap showing the mRNA expression of genes involved in immune-modulatory and senescence associated genes in differentiating aNPCs at DIF4 upon Mili KD compared to control cells. Expression level represented in Transcript per Million (TPM). (B) Bright field microscopy images (left) and quantification (right) of ß-galactosidase positive cells as percent of total cells upon Mili KD or control, 48 h after induction of spontaneous differentiation. (C) Fluorescence microscopy images (left) and quantification (right) of control or Mili KD aNPCs 48h after spontaneous differentiation, immunostained with anti-BrdU (white) and Ki67 (purple) antibodies. (Right) Percentage of BrdU+ and Ki67-cells over BrdU+ cells. (D) (Top) Representative cell cycle analysis of propidium iodide staining by flow cytometry; (Bottom) Percentage of aNPCs in G0/G1, S and G2/M phases 48h after spontaneous differentiation. (E, F) Relative abundance of cell cycle-dependent genes (E) and involved in cell cycle (F) in Control and Mili KD aNPCs 48 hours (E) or DIF4 (F) after spontaneous differentiation. (G) (left) Scheme of the *in vivo* experiment; (right) *Mili* mRNA expression in Nestin+ sorted cells from young (6 weeks) and old (56 weeks) mice. Data are expressed as mean ± SEM, n = 3 independent, (in A, 2 separate flow cells per sample). t-Student test as post hoc: *p < 0.05, **p < 0.01, ***p< 0.001, **** p < 0.0001. Scale bar, 50 µm.

### The piRNA pathway modulates noncoding RNAs and mRNAs involved in ribosome assembly and translation

Next, we sought to identify piRNA pathway targets in aNPCs lineages (Fig.5). In contrast to germline piRNAs which primarily target TEs, somatic piRNAs have also homology with, or pair by sequence complementarity to noncoding RNAs, such as transfer RNAs (tRNAs) (Keam et al., 2014) and others (Rojas-Ríos and Simonelig, 2018). We searched for noncoding RNAs that are putative targets of the piRNAs expressed in our model (Fig. 2). Indeed, TEs were just a minor percentage of the predicted noncoding RNA targets in both undifferentiated aNPCs and progeny (Fig.5 A, B), despite their proportion being increased upon induction of neurogenesis (Fig. 5 B). The latter finding is in agreement with the observed activation of TEs (*i*.*e*., LINE1) during neuronal differentiation (Coufal et al., 2009; Muotri et al., 2005). Interestingly, 5S ribosomal RNA (5S rRNA) and tRNAs were the main predicted piRNA targets in both undifferentiated (47% and 40%, respectively) and progeny (35% and 16%, respectively, Fig. 5 A, B). To ascertain whether these noncoding RNAs are modulated by the piRNA pathway, we quantified their expression in aNPCs and progeny, upon Mili-KD (Fig. 5 C, D). Indeed, piRNA pathway depletion significantly elevated levels of 5S rRNA and SINE-B1 family of TEs in both undifferentiated aNPCs and progeny, compared to scrambled control (Fig.5 C), whereas LINE1, here quantified with a qPCR assay detecting the full-length transcript, was initially refractory to piRNA depletion and its level only increased late in differentiation (Fig. 5 D). These results indicate that 5s rRNAs and SINE-B1 are repressed by the piRNA pathway in neurogenesis. To identify which mRNAs are modulated by the piRNA pathway in neurogenesis, we analyzed mRNA transcriptome by RNA seq from Mili KD or scrambled control cells during their spontaneous differentiation at DIF 4, *i*.*e*., when the piRNAs are most abundant (Fig. 2). Most of the modulated genes upon piRNA pathway depletion were upregulated (Fig 5 E), and the majority of them bore sequences antisense to a piRNA (*i*.*e*., were predicted piRNA targets), or harbored homologous sequences to piRNAs (Fig. 5 F). We found that mRNA transcripts from individual genes are targeted by multiple piRNAs, with a maximum of 11,904 unique piRNA sequences targeting a modulated gene (Fig. 5G). Spearman analysis indicated a positive correlation between piRNA levels in undifferentiated and differentiating aNPCs/neuroblasts, and the degree of target upregulation upon piRNA pathway depletion (Fig. 5H). Gene ontology (GO) analysis of the upregulated piRNA targets in Mili KD cells at DIF 4 indicated a prevalence of “ mRNA processing” and “ Translation” among the top ten biological processes affected (Fig. 5 I). Moreover “ Ribosome” (p-value 7,7 E-8) and “ Splieceosome” (p-value 3,3 E-7) were the top enriched pathways based on function in the Kyoto Encyclopedia of Genes and Genomes (KEGG) database (*not shown*). This evidence is consistent with our observation that 5S rRNA, SINEB1 and possibly tRNAs are modulated by the piRNA pathway in these cells.

**Fig 5.**
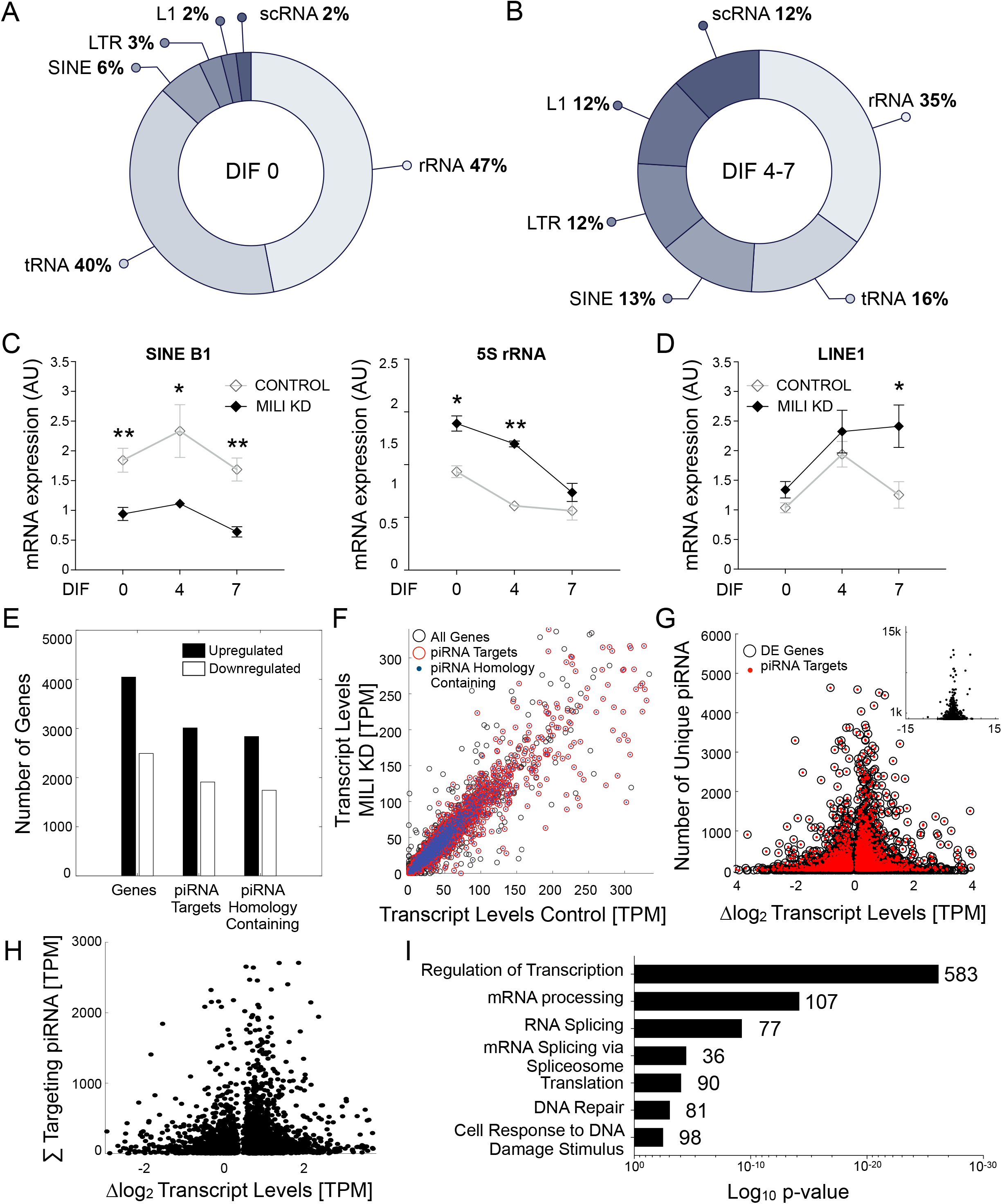
piRNAs target noncoding RNAs and mRNAs involved in ribosome assembly and translation. (A, B) Pie plots showing proportions of noncoding RNAs predicted targets of piRNAs in undifferentiated aNPCs (A, DIF0) and upon induction of neurogenesis (B, DIF4-7). (C) Transcription levels of SINE B1 and 5S rRNA, and (D) transcription levels of LINE1 5’UTR in the Mili KD aNSCs (DIF0) and differentiating aNPC (DIF4-7). (E) Total counts of upregulated (black bars) and downregulated genes (white bars), for all genes, all piRNA targeted genes and those with piRNA homology (indicated). (F) Scatter plot of all detected genes in control or Mili KD aNPCs (black circles), piRNA target (red circles) and those with piRNA homology (blue dots). (G) The log2 fold-change of significantly altered genic mRNA transcripts (abscissa) plotted with the raw number of unique piRNA sequences qualified as targeting molecules (ordinate) for all modulated genes (black circles), and piRNA-targeted genes (red dots), identified by RNA-Seq. The modulated genes without targeting piRNAs concatenate at the bottom at the ‘y = 0’ value. Total rage, inset. (H) The log2 fold-change of significantly altered piRNA-targeted gene mRNA transcripts (abscissa) plotted with the summed levels of all mRNA-targeting (complementary) piRNA molecules (ordinate), identified by RNA-Seq. (I) Bar graph showing the top biological pathways significantly upregulated upon Mili KD; Numbers in each category indicate gene counts. Data are expressed as mean ± SEM, n = 3 independent experiments. t-Student test as post hoc: *p < 0.05, **p < 0.01 (C &D). Significant difference was detected by the one-way ANOVA, p<0.05 (G & H). Data are expressed in transcripts per million (TPM) as the mean levels of 6 sequencing runs for 3 samples. (E-I) transcriptome profiling at DIF4. Outliers (mean calculation) were detected by more than three mean absolute deviations, for a final ‘n’ value between 4 and 6 samples.

### Inhibition of the piRNA pathway in aNPCs enhances polysome assembly and results in higher protein synthesis upon differentiation

As dysregulation of ribosome biogenesis and translation have been associated to cellular senescence (Liu and Sabatini, 2020), we analyzed polyribosomes in aNPCs and in their progeny. To this end we first used stimulated emission depletion (STED) nanoscopy, a method that allows to visualize and quantify polysome density (Viero et al., 2015). Depletion of the piRNA pathway increased polyribosomes in both undifferentiated and differentiating aNPCs, as revealed by immunostaining for the ribosomal protein RPL26, compared to control cells (Fig. 6A, B). As the density of ribosomes over transcript does not necessarily correlate with its translation (Mills and Green, 2017), we also quantified protein synthesis rate by OPP (O-propargyl-puromycin) labelling of nascent proteins during neurogenesis. Accordingly, protein synthesis rate was significantly increased upon piRNA pathway depletion in differentiating cells, but not in undifferentiated aNPCs (Fig. 6C, DIF 7).

**Fig. 6.**
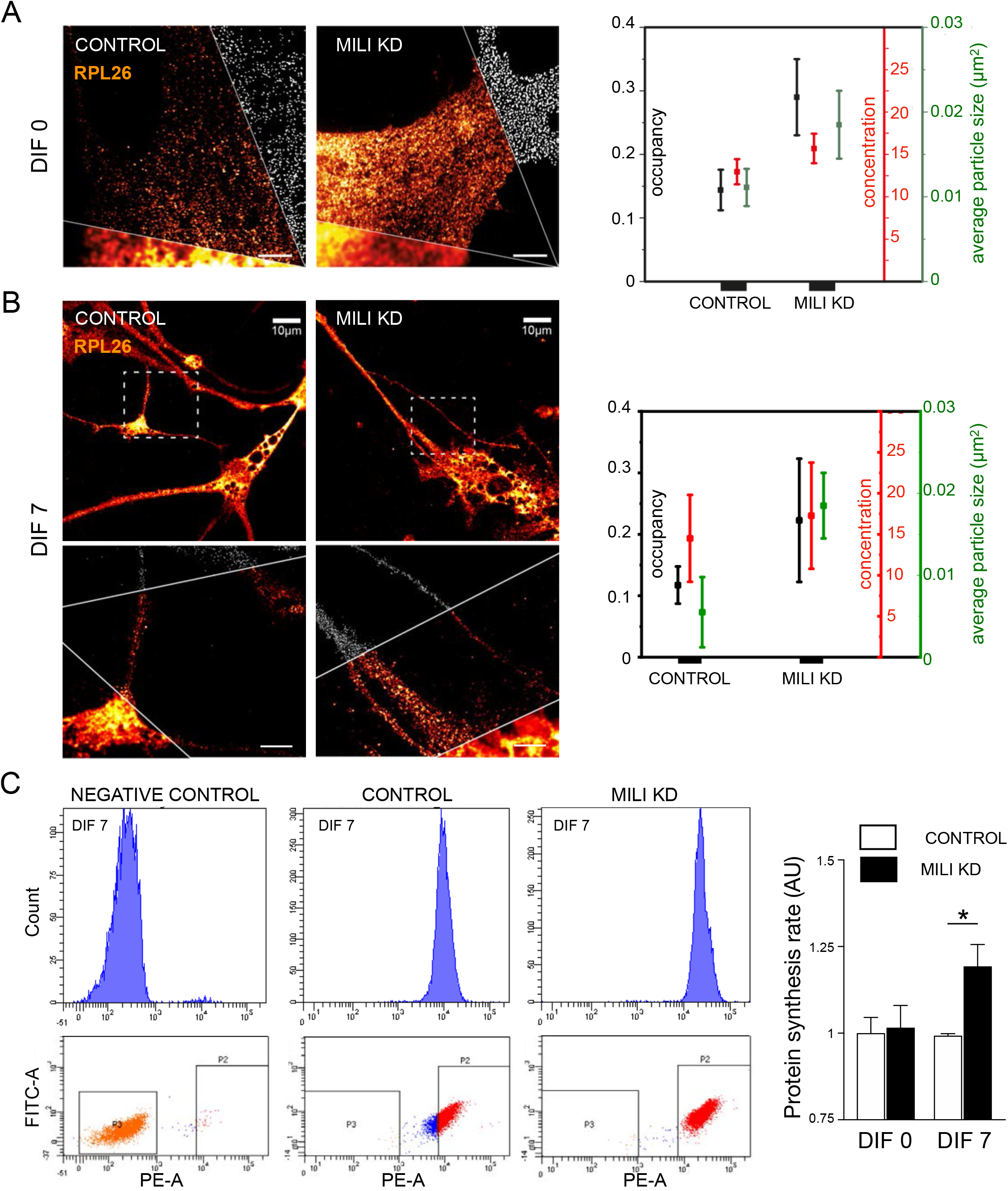
Inhibition of the piRNA pathway in aNPCs enhances polysome assembly and results in higher protein synthesis upon differentiation. (A, B) Microscopy images (Middle cut: g-STED nanoscopy; Bottom: Confocal; Top: analysis) of control and Mili KD aNPCs (DIF 0 and 7) immunostained for the ribosomal protein RPL26. (Right) normalized distributions of the occupancy, concentration and average of particle size of each polyribosome particle in the indicated cells. (C) Protein synthesis rate (right) as determined by OPP incorporation assay with flow cytometry (left) in control and Mili KD undifferentiated (DIF0) and differentiating (DIF 7) aNPCs. Scale bars: 2 (A); 10 (B) μm. Data are expressed as mean ± SEM, n = 3 independent experiments. t-Student test as post hoc: *p < 0.05.

In sum, these results support a role for the piRNA pathway in sustaining adult neurogenesis by repressing translation and senescence in aNPCs.

## Discussion

In this study, we investigated the expression of Mili and Mili-dependent piRNAs in mouse and human NPCs, and inferred functions of this pathway in the regulation of neurogenesis in the adult mouse hippocampus. Our results provide evidence of an essential role for the piRNA pathway in mammalian neurogenesis. This finding adds a new layer of complexity to the understanding of adult brain plasticity and aging, and entails crucial implications for neuronal disorders where dysregulated expression of the piRNA pathway has been reported, such as neurodegeneration (Jain et al., 2019; Wakisaka, 2019).

Differently to the canonical functions of the piRNA pathway in germline, which are mainly required for maintenance of stem cell pools, here we provide the first evidence of a requisite role linking the piRNA pathway with proper differentiation and fate choice of neural stem cells. Specifically, we show that piRNA pathway inhibition in aNPCs leads to aberrant neurogenesis, and to an increased generation of reactive astrocytes, a phenotype which has been observed in the hippocampi of aged mice (Bonaguidi et al., 2011; Clarke et al., 2018; Encinas et al., 2011; Martín-Suárez et al., 2019). Accordingly, we find that inhibition of piRNA biogenesis in aNPCs results in senescence-like phenotype, thus providing a possible cellular mechanism that drives aNPC fate adrift toward reactive astrocytes.

Moreover, and in agreement with previous observations linking cell senescence to altered ribosome biogenesis (Liu and Sabatini, 2020), our data suggest pleiotropic functions of the Mili and Mili-dependent piRNAs in aNPCs reduce polysome assembly and protein synthesis. This result differs from previous functional evidence for the piRNA pathway in germline stem cells, where Miwi/Maelstrom interacting piRNAs activate translation to sustain spermiogenesis (Castañeda et al., 2014; Dai et al., 2019). Despite the precise cascade of events that lead to dysregulation of NSCs homeostasis in aging is only beginning to emerge, proper control of proteostasis, pro-neurogenic, cell cycle and pro-inflammatory signaling is involved in maintenance of lifelong neurogenesis. Our study about Mili and its targets in mouse NSCs certainly provides an entry point to dissect the role of the piRNA pathway in mammalian neurogenesis in the context of brain aging.

## Methods

### Lead contact and materials availability

Further information and requests for resources and reagents should be directed to and will be fulfilled by the Lead Contact, Davide De Pietri Tonelli.

### Materials and Methods

#### Mice

C57BL/6 and Td-Tomato^flox/wt^ knock-in reporter mice (Jackson Laboratory stock number 007908) (Madisen et al., 2010), were housed at *Istituto Italiano di Tecnologia* (IIT); Nestin-GFP mice (Mignone et al., 2004) for the Kainic Acid experiments were housed at the Swammerdam Institute for Life Sciences, University of Amsterdam. All animal procedures were approved by IIT animal use committee and the Italian Ministry of health, or by the Commission for Animal Welfare at University of Amsterdam (DEC protocol 4925, AVD1110020184925), respectively and conducted in accordance with the Guide for the Care and Use of Laboratory Animals of the European Community Council Directives. All mice were group-housed under a 12-h light-dark cycle in a temperature and humidity-controlled environment with ad libitum access to food and water.

### aNPCs cultures

Hippocampal NPCs were prepared and expanded as described previously (Babu et al., 2011; Pons-Espinal et al., 2019; Walker and Kempermann, 2014) and induction of spontaneous differentiation by growth factor removal was done as previously described (Pons-Espinal 2017, and Pons-Espinal, Gasperini et al 2019); viral induced-neuronal differentiation of aNPCs was done by transduction of a viral construct expressing Ascl1-ERT2 as previously described (Braun et al., 2013).

### Protein extraction and Western blot

For total protein extraction, adult testes or hippocampus or cell pellets were homogenized in RIPA buffer and the protein concentration was determined using a Bradford Assay kit (Bio-Rad). For blot analysis, equal amounts of protein (30 µg) were run on homemade 10% polyacrylamide gels and transferred on nitrocellulose membranes (GE Healthcare). Membranes were probed with primary antibodies (listed in the table below), followed by HRP-conjugated secondary antibody anti-rabbit or mouse (Invitrogen, A16104, A16072; 1:2,000). The band corresponding to Mili protein was detected with two different antibodies and validated with four different RNAi constructs (1 shRNA virus and 3 GapmeRs). LAS 4000 Mini Imaging System (GE Healthcare) was used to digitally acquire chemiluminescence signals, and the band intensities were quantified using Fiji (Macbiophotonics, Fiji is Just ImageJ) (Schindelin et al., 2012). List of antibodies used available in the SI appendix.

### Virus and GapmeR injection

Virus or GapmeR injection was done as previously shown (Pons-Espinal et al., 2019): 8 weeks-old mice Td-Tomato^flox/wt^ or WT C57BL6/J were anesthetized with isoflurane, 1 μl of virus mix (Split-Cre N-Cre:C-Cre) or 1.5 μl of antisense LNA GapmeR for Mili-KD or negative control (MILI 339511, Control 339515, Qiagen), were stereotaxically injected in the dentate gyrus. To assess the efficacy of *Mili* knockdown, a first group of mice (n=5) was sacrificed 48 hours after the injection and the DG dissected for RNA or protein extraction. 24 hours after the oligos injection another set of animals received 2 BrdU intraperitoneal injections per day for 5 days (50 mg/kg) (one every 12 hours). Animals were sacrificed 10 (n=5) or 30 days after oligos injection (n=7) for histological analysis, as previously described (Pons-Espinal et al., 2019). Mice were anesthetized with intraperitoneal administration of ketamine (90mg/kg) and xylazine (5-7mg/kg), and subsequently perfused with PBS followed by 4% paraformaldehyde (PFA). Brains were harvested, postfixed overnight in 4% PFA, and then equilibrated in 30% sucrose. 40 μm brain sections were generated using a sliding microtome and were stored in a −20°C freezer as floating sections in 48 well plates filled with cryoprotectant solution (glycerol, ethylene glycol, and 0.2 M phosphate buffer, pH 7.4, 1:1:2 by volume). Slices were used for immunofluorescence and immunohistochemical analysis.

### Fluorescence-Activated Cell Sorting (FACS) and flow cytofluorometric analysis of cell cycle distribution

For RNA extraction and cDNA preparation, Td-Tomato^flox/wt^ or Nestin-GFP mice were used. Six to ten Td-Tomato^flox/wt^ mice were euthanized 10 or 30 days after the split cre viruses injection. DG cells were dissociated with the Neural Tissue Dissociation Kit P (Miltenyi Biotec) and FACS-sorted as previously published (Walker et al., 2016). FACS-sorted cells were immediately processed for RNA extraction. To measure the cell cycle length, we used propidium iodide (PI), which binds to DNA by intercalating between the bases, as previously described (Krishan). Briefly, cells were trypsinized, resuspended in PBS and fixed with 70% of ethanol for 40 minutes on ice. Cells were then centrifuged, resuspended in PBS for 15 minutes and then incubated 1h at 37°C with 60 µg/ml of PI (Sigma). Cells were collected by centrifuge and resuspended in ice-cold PBS for FACS analysis.

### Immunostaining analysis

The immunostaining on brain slices was performed on sections covering the entire dorsal hippocampus as previously described (Paxinos, 2001; Pons-Espinal et al., 2019). To detect Ki67 staining, citrate buffer 10 mM pH = 6 treatment during 10 min at 95 °C was used. Primary antibodies are listed below, secondary fluorescent antibodies were diluted 1:1000 (Goat Alexa 488, 568, and 647nm, Invitrogen). Confocal stack images of brain slices (40µm) were obtained with the Confocal A1 Nikon Inverted SFC with 20x objective. Cell quantification and analysis was performed using NIS-Elements software (Nikon) and the Cell-counter plugin in Fiji. Immunofluorescence staining on cell cultures was performed as reported in (Pons-Espinal et al., 2019). To detect BrdU incorporation, cells were pretreated with 2N HCl for 30 min at 37°C. Cells were mounted in mounting medium and counterstained with fluorescent nuclear dye DAPI (Invitrogen). Images were obtained using the microscope Nikon Eclipse at 20x or 40x magnification and quantification was performed using a Cell-counter plugin in Fiji. DAB staining was performed as previously reported (Bielefeld et al., 2019). Briefly, sections were incubated with peroxidase block (Vectashield), permeabilized with 0.3% PBS-Triton X (PBS-T) and 0.1% PBS-T. Sections were blocked with 0.1% PBS-T and 5% Normal Goat Serum (NGS), incubated with primary antibodies and subsequently with the corresponding biotinylated secondary antibodies (1:1000 Goat anti-rabbit, Invitrogen). Signal amplification was performed using the ABC complex (Vectashield), according to manufacturer’s instructions. Sections were incubated with the solution for DAB reaction (Sigma) and counterstained with Hoechst (1:300), mounted and cover slipped with Vectashield reagent (VECTOR Labs). List of antibodies used available in the SI Appendix.

### RNA extraction and real time qPCR

Total RNA was extracted from aNPCs (proliferating and differentiating conditions), or DG dissected from adult C57BL/6, Nestin-GFP or Td-Tomato^flox/wt^ mice with QIAzol protocol (Qiagen) according to the manufacturer’s instructions. 1 µg of total RNA was treated with DNase I (Sigma) and cDNA was synthesized using iScript cDNA Synthesis kit (Bio-Rad) or with ImProm-II reverse transcriptase (Promega). Real time qPCR was performed in duplex with Actin or Ubiquitin C as a reference gene, with QuantiFast SYBR Green PCR Kit (Qiagen) or Taqman Assay (Thermo Fisher) on ABI-7500 Real-Time PCR System (Applied Biosystems). Expression levels were determined relative to Ubiquitin C or Actin, using delta delta Ct method. Primers listed below were designed using NCBI/UCSC Genome Browser and Primer3 software tools and then checked in PrimerBLAST for their specificity to amplify the desired genes. List of primers used available in the SI Appendix.

### Small RNA library preparation

Cells were lysed in QIAzol lysis reagent and total RNA was isolated using the miRNeasy Mini kit (Qiagen), according to the manufacturer’s instructions. Quantity and quality of the total RNA were measured by Nanodrop spectrophotometer and Experion RNA chips (Bio-Rad). RNA with RNA integrity number (RIN) values ≥ 9.5 were selected for the study. 1 μg of high-quality RNA for each sample was used for library preparation according to the Illumina TruSeq small RNA library protocol (Illumina Inc., CA). Briefly, 3’ adapters were ligated to 3’ end of small RNAs using a truncated RNA ligase enzyme followed by 5’ adaptor ligation using RNA ligase enzyme. Reverse transcription followed by PCR was used to prepare cDNA using primers specific for the 3’ and 5’ adapters. The amplification of those fragments having adapter molecules on both ends was carried out with 13 PCR cycles. The amplified libraries were pooled together and run on a 6% polyacrylamide gel. The 145-160 bp bands (which correspond to inserts of 24-32 nt cDNAs) were extracted and purified using the Wizard® SV Gel and PCR Clean-Up System (Promega). The quality of the library was assessed by the Experion DNA 1K chips (Bio-Rad). Small RNA sequencing using HiSeq2000 (Illumina Inc., CA) was performed by the Center for Genomic Science of IIT@SEMM.

### Mili knock down *in vitro*

To constitutively knockdown Mili expression *in vitro*, aNPCs were infected at MOI=5 with a lentivirus encoding for a *Mili-*targeted short hairpin (shMILI, plKO.1, Sigma), or for a short hairpin scramble lentivirus (Control, SHC202, Sigma) both decorated with an eGFP reporter. GFP-positive cells were first selected by FACS after three passages, and then plated in proliferating or differentiating media, as previously described. We also performed the knock down using two different synthetic antisense LNA GapmeRs for Mili-KD or negative control (Mili 339511, Control 339515, Qiagen). Mili knock down was assessed by real time qPCR and Western Blot.

### mRNA library preparation

aNPCs WT and transduced with the Control or with the shMILI lentiviruses, were grown to confluence and allowed to spontaneously differentiate for the indicated days. Total RNA was extracted using QIAzol lysis reagent and quantity and quality of the total RNA were measured by Qubit 4 Fluorometer (Thermo Fisher) and Bioanalyzer RNA chips (Agilent). RNA with RNA integrity number (RIN) values ≥ 8 were selected for the study. 30 ng of high-quality RNA for each sample was used for library preparation according to the Illumina Stranded Total RNA Prep, Ligation with Ribo-Zero Plus Kit (20040529, Illumina Inc., CA) using the IDT for Illumina Indexes Set A (20040553). Briefly, after ribosomal RNA depletion, RNA was fragmented and denatured and cDNA synthesized. Then the 3’ ends were adenylated, and anchors ligated. After amplification and clean up, quality of the libraries was assessed by the Bioanalyzer DNA chips (Agilent). Paired-End stranded total RNA sequencing on NovaSeq 6000 Sequencing System instrument (Illumina Inc., CA), was performed by the IIT Genomic Unit @Center for Human Technologies.

### Small RNA sequencing data processing

Illumina reads were trimmed to remove the 3′ adapter using Cutadapt, with parameters -m 25 -q 20. Since piRNA size ranges from 26 to 31 bases, all sequences with length ≤24 bases were discarded. Reads mapped to known non-coding RNAs (RNAcentral v6.0 snoRNA, UCSC tRNA, miRBase Release 21 miRNA hairpin and mature miRNA annotation, NCBI complete ribosomal DNA unit) (Karolchik, 2004; Kozomara and Griffiths-Jones, 2014; Petrov et al., 2015) were removed from the datasets. The comparison was performed using NCBI BLASTN v2.6.0 with parameters -max_hsps= 1, -max_target_seqs= 1, -perc_identity= 80, mismatches <= 1, qcovhsp >= 90 (Altschul et al., 1990). Reads were aligned on the non-repeat-masked UCSC release 9 of the mouse genome (MM9) (Waterston et al., 2002) using bowtie2 (Langmead and Salzberg, 2012) v2.2.6 with the sensitive preset option and allowed a maximum of 100 alignments. All the reads that aligned to the genome were retained and used for subsequent analysis. piRNA clusters were identified collapsing overlapped piRNA sequences (piRBase Release 1) (Zhang et al., 2014) into one cluster (mergeBed with preset options) (Quinlan and Hall, 2010). piRNA clusters and all the reads that aligned to the genome were intersected (intersectBed with option -f 1). Intersection files were then parsed using a custom perl script in order to evaluate alignment counts. Differential expression was assessed using DESeq2 (Love et al., 2014). piRNA clusters were considered differentially expressed when the adjusted p-value was ≤ 0.05, and down-and up-regulation was established in the range of ≤ -1 to ≥ 1 log2 fold-change, respectively. piRNA sequences were then categorized for the putative mRNA transcript targets (for each gene). In order to obtain a count of piRNA target levels which target individual gene transcripts, for each differentiation time-point (DIF0-7), piRNA transcripts were expressed in transcripts per million (TPM), and summed for each target category. Spearman correlation was performed between the levels of the piRNA in the Sh-Scramble (control) genotype, and compared to the fold-change level of the putative target genes which were found to be significantly altered (up and down) in the Sh-Mili-KD genotype. In order to assess the clustering behavior of putative piRNAs, the 5’ termini positions of each cluster-associated putative primary and putative secondary piRNA sequences were analyzed for distance, represented as probability within a range of 200 nucleotides in the 5’ direction and 200 nucleotides in the 3’ direction of the putative primary piRNAs, as reported previously (Gainetdinov et al., 2018). The positional distance between piRNAs for each cluster was sampled iteratively and normalized by the total number of diverse piRNAs associated with each cluster. The distance probability distribution was assayed by the locally weighted smoothing linear regression method (LOWESS), by using the built-in MATLAB ‘fit’ function (MathWorks, Natick, MA), with a span value of 0.1.

### mRNA Sequencing data processing

Illumina sequencing was performed bidirectionally, and in duplicate by two flowcell pairs of 100 and 150 base pairs, for a total of 6 measurements produced from 3 independent samples for each differentiation time-point and genotype. Adapter sequences were trimmed using *Cutadapt*, after which a quality control trim was implemented on a sliding window of 25 nucleotides, for 2 base pairs with a minimum quality score of 26, with the *Bowtie* build for the MATLAB bioinformatics suite (MathWorks, Natick, MA). Transcript quantification was performed with the *Salmon* suite, as reported previously (Patro et al., 2017), on the NCBI mouse genome, release 67, obtained from the ENSEMBL FASTA directory. The identified ENSEMBL gene accessions were grouped for the different transcript reads, and the read counts, expressed in TPM, were summed for the annotated transcripts. Outlier reads for each gene transcript level in TPM were detected by the mean absolute deviations method (MAD), where reads with more than 3 scaled MAD distances from the mean were eliminated from statistical analysis. Then, the mean and SD for each gene were used for statistical analysis among the different genotypes by one-way ANOVA with multiple comparison. *P*-values lower than, or equal to, 0.05 were selected as the threshold of significance, for a minimum count of 4 of 6 samples per gene. Database for Annotation, Visualization and Integrated Discovery (DAVID, https://david.ncifcrf.gov/), (Huang et al., 2009), was used to perform Gene ontology (GO) and Kyoto Encyclopedia of Genes and Genomes (KEGG) signaling pathway analysis.

### Periodate oxidation and alkaline ß-elimination

Periodate oxidation and alkaline ß-elimination was performed as previously described (Balaratnam et al., 2018). Briefly, total small RNAs were isolated using miRneasy Mini kit (Qiagen) according to the manufacture’s protocol. Each sample was split in two portions, each containing 25µg of RNA and independently treated with either 200 mM of sodium periodate or water, and RNA precipitated by ethanol. As internal control for the assay, a synthetic piRNA sequence (corresponding to one of the most abundant piRNA found in aNPCs – piR-cluster-1) either modified with the 3’-end 2’-O-methylation (positive control) or bearing a terminal 2’,3’-hydroxyl group (negative control) were subject to the periodate treatment as above. The samples and control oligos were finally quantified by qPCR-TaqMan small RNA assay.

### piRNAs real time qPCR

Total RNA enriched in the fraction of small RNAs, was extracted using miRNeasy Mini kit following the manufacturer’s instructions from aNPCs, DG extracted from C57BL6/J or Td-Tomato^flox/wt^ mice. cDNA was obtained using the TaqMan MicroRNA Reverse Transcription Kit (Thermo Fisher) according to the manufacturer’s instructions and quantified using the Custom TaqMan Small RNA Assay (Thermo Fisher) on a ABI-7500 Real-Time PCR System (Applied Biosystems). Each sample was normalized to U6 snRNA level (Thermo Fisher). Cluster sequences used for probe design are listed in Table S1.

### ß-galactosidase detection

Was obtained with the Senescence Cells Histochemical Staining Kit (Sigma-Aldrich, CS0030), according the manufacturer’s instructions. Briefly, cells were plated on coverslip in proliferating medium. 48h after the induction of spontaneous differentiation, cells were washed twice with PBS 1X and incubated with Fixation Buffer 1X for 7 min at RT. Next, cells were rinsed with PBS 1X and incubated with fresh senescence-associated ß-Gal (SA-ß-Gal) stain solution at 37°C (no CO2) for 4 h. Reaction was blocked with PBS 1X and coverslips were mounted on slides using Vectashield reagent (VECTOR Labs). Images were obtained using the microscope Nikon Eclipse, the percentage of cells expressing ß-galactosidase was quantified over the number of total cells using a Cell-counter plugin in Fiji software.

### *in silico* piRNA targets prediction

For piRNA targets analysis, we divided the sequencing data in one set of 100 piRNA clusters enriched in proliferating aNPCs (DIF0) and a second set of 198 clusters specifically expressed at DIF4/7 stage. The Differential Expression analysis for piRNAs mapping on REs showing trends for enrichment of piRNAs mapping on repeat elements (REs) in DIF4 and DIF7 compared to DIF0 was done using EdgeR software package (Robinson and Oshlack, 2010). Identification of piRNA targets was divided in: piRNAs mapping on REs only / piRNAs mapping on GENCODE elements / piRNAs mapping on REs within GENCODE elements / unannotated piRNAs / piRNAs clusters. Gene Ontology analysis for piRNAs mapping on GENCODE protein-coding genes (but NOT mapping on REs) has been done with the R package GOFuncR (https://bioconductor.org/packages/release/bioc/html/GOfuncR.html)

### Protein synthesis assay

To quantify the protein synthesis rate of cells, we used the Global Protein Synthesis Assay Kit (FACS/Microscopy), Red Fluorescence kit (abcam), following the manufacturer’s instructions. Briefly, cells in proliferating or differentiating media (DIF7) were treated with Cycloheximide as an inhibitor of protein synthesis, for 30 min at 37°C. Media were replaced with fresh aliquots containing Protein Label (400X) diluted to 1X final concentration and the cells were incubated for additional 30 min at 37°C. Negative control cells were not incubated with the protein label. Samples were analyzed by FACS for red fluorescence generated by *de novo* synthesized protein during click reaction. Translation rate is directly proportional to emitted fluorescence. Cells emitting fluorescence lower than 10^3^ were considered negative (P3), higher than 10^4^ were considered positive (P2).

### Kainic Acid Administration, single cell suspension and enrichment of aNPCs by FACS

Kainic Acid to elicit tonic, non-convulsive epileptic seizures, was administered as described before (Bielefeld et al., 2019). Briefly, 50nL of 2.22mM Kainic Acid dissolved in PBS (pH 7.4) was injected bilaterally into the hippocampus at the following coordinates (AP -2.0, ML +/-1.5, DV -2.0 mm) (between 9AM and 1PM). Control animals were administered saline (pH 7.4). Bilateral dentate gyri from 3 animals per condition were pooled to allow sufficient recovery of NPCs. A single cell suspension was created using a Neural Tissue Dissociation kit (Miltenyi Biotec), according to the manufacturers protocol. In order to enrich aNPCs from the DG, we used the endogenous GFP expression driven by the Nestin promotor in combination with FACS. Propium Iodide (5µg/mL) was added to the single cell suspension to assess cell viability. Cells were sorted using a FACSAriatm III system (BD) with 488nm excitation laser. Cell duplets were removed based on forward and side scatter and viable cells were selected based on PI negativity. GFP-positive (corrected for autofluorescence) cells were sorted (≅ 50000 cells/pool) and collected in PBS containing 1% FBS. Trizol LS (Thermo Scientific) was added and after resuspension samples were snap-frozen and stored at -20°C.

### Immunofluorescence, STED nanoscopy and particle analysis

Confocal and Stimulated Emission Depletion (STED) nanoscopy were performed as previously reported (Diaspro and Bianchini, 2020; Vicidomini et al., 2018). aNPCs were plated on glass coverslips 24 h before fixation. Cells were fixed with PFA 4%, permeabilized with PBS-T 0.1%, blocked 1 hour at room temperature with PBS-T 0.1% NGS 5% and incubated according to the dilution suggested by the manufacturer’s instructions with 0.01 μg/ml rabbit polyclonal antibody against the N terminus of RPL26 (Abcam) for 1 h at room temperature. Cells were washed extensively and incubated with the secondary antibody goat anti–rabbit ATTO-647N (0.8 μg/ml; Sigma) for 45 min. Nuclei were stained while mounting the coverslip with DAPI-Prolong antifade (Invitrogen). Confocal and STED images were acquired at 23°C with a modified TCS SP5 STED-CW gated (Leica Microsystems, Mannheim, Germany) operated with Leica’s microscope imaging software. The microscope has been customized with a second pulsed STED laser line at 775nm. The beam originates from a Onefive Katana HP 8 (NKT, Birkerød, Denmark) and pass through a vortex phase plate (RPC photonics, Rochester, NY, USA) before entering the microscope through the IR port. The depletion laser pulses are electronically synchronized with the Leica’s supercontinuum pulsed and visible excitation laser. The ATTO-647N fluorescence was excited at 633 nm, and the fluorescence depletion was performed 775 nm. The maximal focal power of the STED beam was 200 mW at 80 MHz. Both beams were focused into the 1.4 NA objective lens (HCX PL APO 100× 1.40 NA Oil STED Orange; Leica). Fluorescence was collected by the same lens, filtered with a 775 nm notch filter, and imaged in the spectral range 660–710 nm by hybrid detector with a time gating of 1 ns. All of the images have 14-nm pixel size and 37-μs pixel dwell time. We performed the analysis of polysome clusters in aNPCs on more than 20 images likewise different cells. Image analysis was performed using the Fiji software.

### Quantification and Statistical analysis

Data are presented as mean ± SEM and were analyzed using Prism 6 (GraphPad). Statistical significance was assessed with a two-tailed unpaired t test for two experimental groups. For experiments with three or more groups, one-way ANOVA with Bonferroni’s multiple comparison test as post hoc was used. Results were considered significant when p < 0.05.

## Data and code availability

RNA sequencing data (mouse) have been deposited in the European Nucleotide Archive (ENA) at EMBL-EBI and available upon request; human datasets are available through RIKEN FANTOM5.

## Acknowledgments

We are grateful to G. Hannon (Cambridge, UK) for providing Mili, Miwi antibodies; G. Enikolopov (Stony Brook Univ. NY, USA) for the Nestin-GFP mouse. We thank IIT technical staff (S. Bianchi; M. Pesce; E. Albanesi, M. Morini, D. Vozzi) for excellent assistance and L. Pandolfini (IIT-CHT Genoa, Italy) for advice in piRNA resistance to periodate oxidation and alkaline ß-elimination protocol.

## Author contributions

Conceptualization: C.G., D.D.P.T; Methodology and experiments: C.G, K.T., R.P., D.M., P.B. (UVA), C.P.F., P.B. (IIT), M.S., M.P.E.; *In silico* analysis: A.L.V, R.M.C., G.P., K.T.; Data Curation and visualization: C.G.; Writing – original draft preparation: C.G., D.D.P.T; –Editing: all the authors; Supervision: R.S., A.D., C.P.F., P.C., S.G.; D.D.P.T.; Project administration: D.D.P.T.; Funding acquisition: D.D.P.T.

