## Supplementary materials and figures for "The piRNA pathway sustains adult neurogenesis by reducing protein synthesis and cellular senescence"

**This file includes:**

- **Supplementary Material**
- **Supplementary Figures S1-S3**
- **Supplementary data Table S1** (legend)
- **Supplementary data Table S2** (legend)

### List of antibodies used for WB:

| Antibody | Host | Company | Catalog | Dilution |
|---|---|---|---|---|
| MILI | Mouse | Santa Cruz | sc-377347 | 1:100 |
| | Rabbit | Kind gift of Dr. Hannon | | 1:150 |
| MIWI | Rabbit | Kind gift of Dr. Hannon | | 1:200 |
| ACTIN | Rabbit | Abcam | ab13970 | 1:1000 |
| GADPH | Rabbit | Santa Cruz | sc-25778 | 1:1000 |
| GFAP | Rabbit | Dako | Z-0334 | 1:1000 |

### List of antibodies used for IF/IHC:

| Antibody | Host | Company | Catalog | Dilution |
|---|---|---|---|---|
| MILI | rabbit | Hannon Lab | | 1:100 |
| BrdU | rat | Abcam | ab6326 | 1:200 |
| KI67 | rabbit | Abcam | ab15580 | 1:250 |
| GFAP | rabbit | Dako | Z-0334 | 1:1000 |
| NeuN | mouse | Millipore | MAB377 | 1:250 |
| RPL26 | rabbit | Abcam | ab59567 | 1:500 |
| Nestin | mouse | Millipore | MAB353 | 1:250 |
| Cleaved Caspase-3 | rabbit | Cell Signaling Technology | 9664 | 1:400 |

### List of Primers used for real time qPCR

| Assay ID | Forward primer (5' to 3') | Reverse primer (5' to 3') |
|---|---|---|
| Actin | GGCTGTATTCCCCTCCATCG | CCAGTTGGTAACAATGCCATGT |
| Mili | GGCCAGCATAAATCTCACAC | TAGCTGGCCATCAGACACTC |
| Miwi | TAATTGGCCTGGAGTCATCC | GAGGTAGTAGAGGCGGTTGG |
| Gfap | GGGGCAAAAGCACCAAAGAAG | GGGACAACTTGTATTGTGAGCC |
| Complement C3 | CCAGCTCCCCATTACGTCTG | GCACTTGCCTCTTTAGGAAGTC |
| Serpina 3n | ATTTGTCCCAATGTCTGCGAA | TGGCTATCTTGGCTATAAAGGGG |

| | | |
|---|---|---|
| Cxcl10 | CCAAGTGCTGCCGTCATTTTC | GGCTGGCAGGGATGATTTCAA |
| Cyclin A | GCCTTCACCATTCATGTGGAT | TTGCTGCGGGTAAAGAGACAG |
| Cyclin D1 | GCGTACCCTGACACCAATCTC | CTCCTCTTCGCACTTCTGCTC |
| Cyclin E | GATCCAGAAAAAGGAAGGCAAA | TGAAGAAATTGCCAAGATTGACA |

| **Assay ID** | **Forward primer (5' to 3')** | **Reverse primer (5' to 3')** | **Probe (5' to 3')** |
|---|---|---|---|
| L1 5'UTR Tf | TGAGCACTGAAACTCAGAGGAG | GATTGTTCTTCTGGTGATTCTGTT A | FAM-GAATCTGTCTCCCAGGTCTG-MGBNFQ |
| SINE B1 | TGGCGCACGCCTTTAATC | GAGACAGGGTTTCTCTGTGTAGC C | FAM-CAGAGGCAGGCGGAT-MGBNFQ |
| 5s rRNA | ACGGCCATACCACCCTGAA | GGTCTCCCATCCAAGTACTAACC A | FAM-CCGAGATCAGACGAGAT-MGBNFQ |
| Ubiquitin C | ACAGACGTACCTTCCTCACC | CCCCATCACACCCAAGAACA | VIC-AAAAAGAGCCCTCCTTGTGC-MGBNFQ |

### Supplementary figures

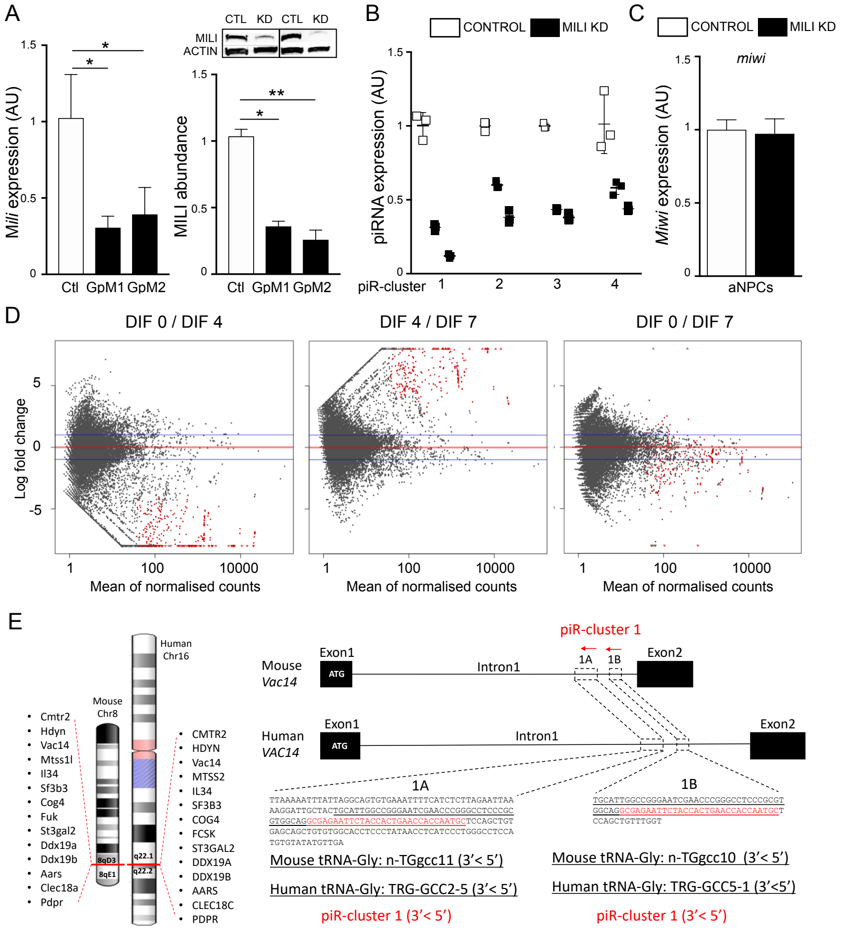


**Fig. S1. KD of Mili does not affect Miwi mRNA levels, and deplete piRNAs abundantly expressed in aNPCs.** Related to Figure 2.

(A) *Mili* mRNA (left) and protein (right) levels in aNPCs upon transfection with control GapmeR (Ctl) or two different GapmeRs (GpM1, GpM2) targeting *Mili* (Mili KD). (B) Expression of four of the most abundant piRNA clusters in control and Mili KD by two different GapmeRs aNPCs. (C) Relative expression of *Miwi* gene in undifferentiated aNPCs in vitro transduced with viruses transcribing a Scrambled short-hairpin (Control) and shMILI (Mili KD). (D) Pairwise comparison of 298 piRNA clusters differentially expressed in undifferentiated aNPCs (DIF0) or upon viral-induced neurogenesis (DIF4-7). (E) Chromosomal location of piR-cluster 1 in mouse and human; (Right) genomic location and sequences (underlined red text) of piR-cluster 1 corresponding to tRNA-Gly genes (underlined black text). Data are expressed as mean ± SEM, n = 3 (A-C) and n = 2 (D-E) independent experiments. t-Student test as post hoc: $*p < 0.05$, $**p < 0.01$.

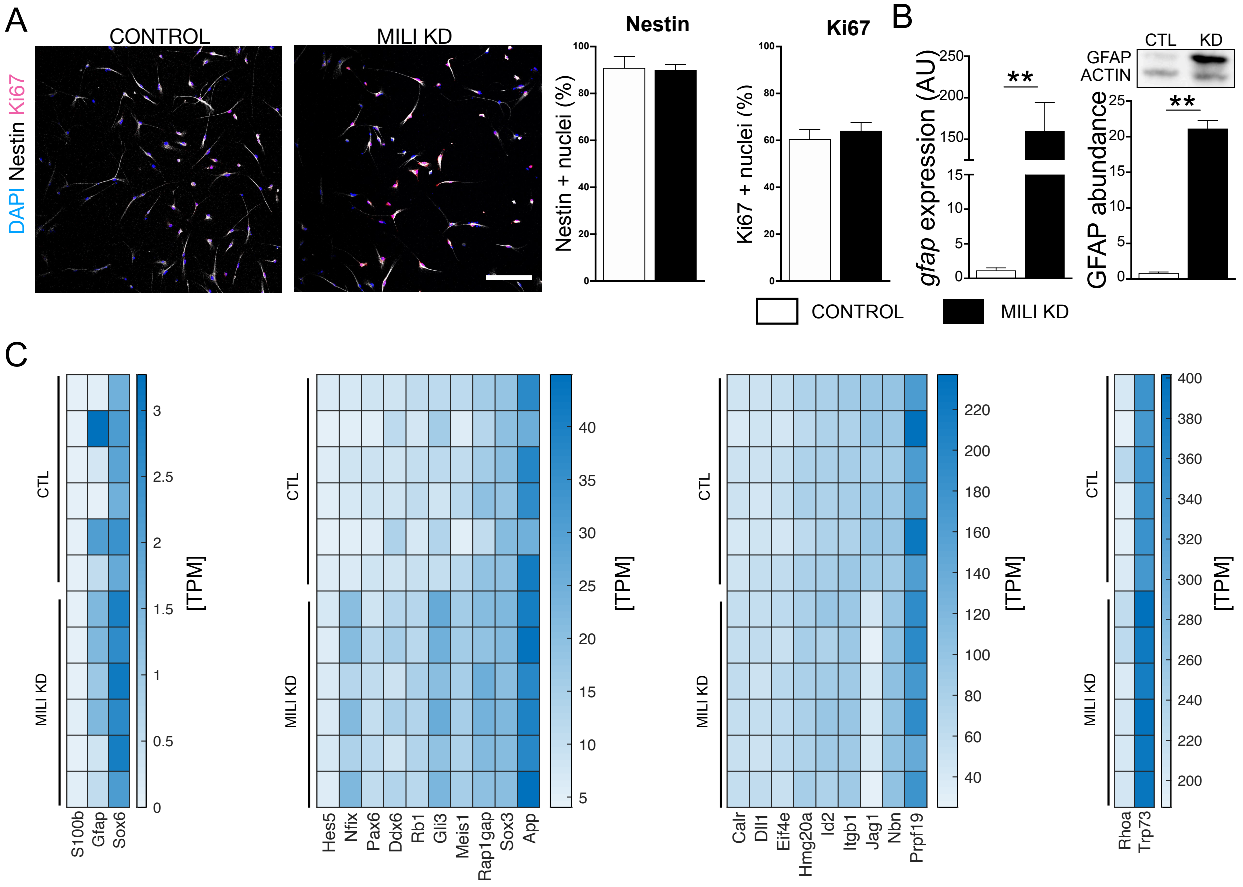


**Fig. S2. Mili KD does not alter aNPC stemness and proliferation, and induces the expression of genes involved in astrogliogenesis.** Related to Figure 3.

(A) (left) Confocal microscopy images of undifferentiated aNPCs *in vitro* transduced with viruses transcribing a Scrambled (Control) and shMILI (Mili KD), immuno-stained with anti-Nestin (white), or anti-Ki67 (purple) antibodies and stained for nuclear DNA with hoechst (blue); (right) Percentage of Nestin or Ki67 positive cells over total cells. (B) *Gfap* mRNA (left) and protein (right) levels in control and Mili KD in differentiated cells (DIF 7). (C) RNA seq. expression data of genes involved in astrocytes development and astrogliogenesis and regulation of neuronal fate in Mili KD aNPCs DIF4 compared to Scrambled control. Data are expressed as mean ± SEM, n = 3 independent experiments, (in C, 2 separate flow cells per sample). **$p < 0.01$. Scale bar 50 µm.

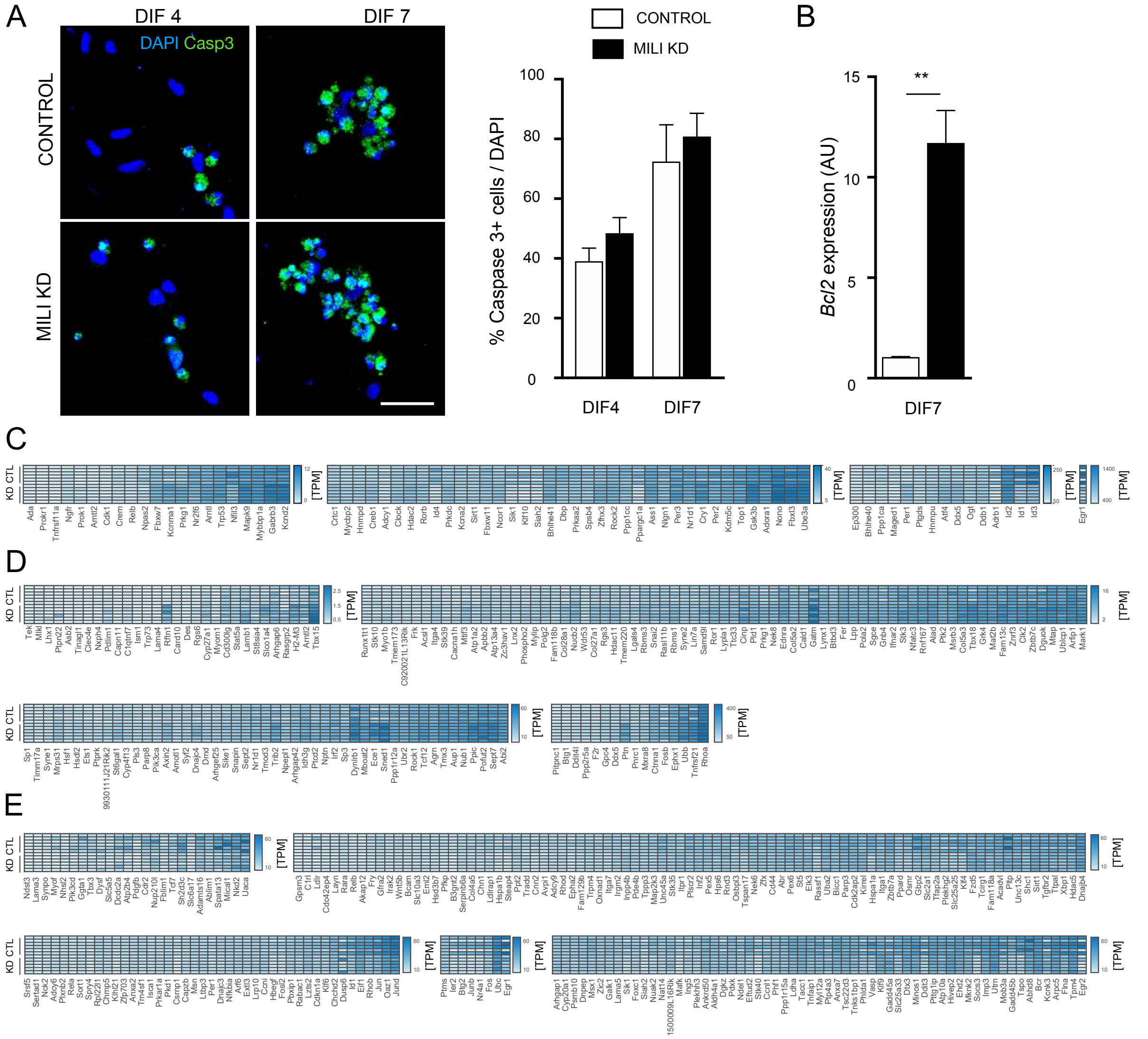


**Fig. S3. piRNA pathway depletion does not lead to apoptosis, rather alters the expression of genes involved in inflammatory responses.** Related to Figure 4.

**(**A) Fluorescence microscopy images of Control or Mili KD cells 4 or 7 days after spontaneous differentiation (DIF 4, 7), immunostained with anti-cleaved caspase-3 (green) and for nuclear DNA with Hoechst (blue). (Right) Percentage of cleaved caspase-3+ cells over total cells. (B) *Bcl2* mRNA expression level in control or Mili KD cells 7 days after induction of spontaneous differentiation (DIF7). (C-E) Heatmap of differentially expressed transcripts in RNA seq from Mili KD neuroblasts, involved in circadian regulation (C, upregulated) or ROS production (D, upregulated; E, downregulated). Data are expressed as mean ± SEM, n = 3 independent (in C-E, 2 separate flow cells per sample). t-Student test as post hoc: $**p < 0.01$. Scale bar, 50 µm.

**Supplementary Data Table S1.**

Genomic location on mouse genome (NCBI37/mm9) and read sequence of piRNA clusters in undifferentiated aNPCs (DIF0) and upon induction of neuronal differentiation (DIF4-7).

**Supplementary Data Table S2.**

Genomic location on human genome (hg38) of piRNA clusters expressed in human neural stem cells (NSC) and astrocytes.
